# The male mouse meiotic cilium emanates from the mother centriole at zygotene prior to centrosome duplication

**DOI:** 10.1101/2022.10.20.512932

**Authors:** P López-Jiménez, S Pérez-Martín, I Hidalgo, FR Garcia-Gonzalo, J Page, R Gómez

**Author notes:** These authors contributed equally to this work (P López-Jiménez, S Pérez-Martín).

## Abstract

Cilia are hair-like projections of the plasma membrane with an inner microtubule skeleton known as axoneme. Motile cilia and flagella beat to displace extracellular fluids, playing important roles in the airways and reproductive system, among others. Instead, primary cilia function as cell type-dependent sensory organelles, detecting chemical, mechanical or optical signals from the extracellular environment. Cilia dysfunction is associated with genetic diseases called ciliopathies, and with some types of cancer.

Cilia have been recently identified in zebrafish gametogenesis as an important regulator of the *bouquet* conformation and recombination. However, there is very little information about the structure and functions of cilia in mammalian meiosis. Here we describe the presence of cilia in male mouse meiotic cells. These solitary cilia form transiently in 20% of zygotene spermatocytes and reach considerable lengths (up to 15 μm). CEP164 and CETN3 localization studies indicate that these cilia emanate from the mother centriole, prior to centrosome duplication. In addition, the study of telomeric TFR2 suggests that these cilia are not directly related to the *bouquet* conformation during early male mouse meiosis. Instead, based on TEX14 labeling of intercellular bridges in spermatocyte cysts, we suggest that mouse meiotic cilia may have sensory roles affecting cyst function during prophase I.

## INTRODUCTION

Centrioles are conserved microtubule-based intracellular structures that form the core of the centrosome and act as templates for the formation of cilia and flagella [1, 2]. Centrosomes are made up of two orthogonally barrel-shaped centrioles embedded in an electron-dense material called the pericentriolar matrix [3]. Each of the centrioles is composed of nine microtubule (MT) triplets around a central proteinaceous structure whose shape resembles a cartwheel [4]. The function of centrosomes in animal cells is absolutely essential, since they behave as the MT organizing centers (MTOCs), controlling the configuration of the MT cytoskeleton of interphasic cells, as well as regulating the formation of the bipolar spindle during cellular division [5-8]. For this reason, centrosome regulation is tightly coordinated with cell cycle progression [2]. In G1, the centrosome is composed of two parental centrioles, but it undergoes a duplication during S phase, with each of the parental centrioles giving rise to a newly formed procentriole. Thus, by G2, cells contain two centrosomes, each consisting of an older and a younger centriole (known as the mother and daughter centrioles, respectively). The two centrosomes then occupy opposite poles of the dividing cell, where they shape the bipolar spindle [9].

Cilia are MT-based plasma membrane projections that can function as motors and/or sensors. Motile cilia beat to displace extracellular fluids, as occurs for example in the mammalian airways, reproductive tract and brain ventricles, where these cilia displace mucus, oocytes and cerebrospinal fluid, respectively [10]. Flagella, despite their different name, are also in essence motile cilia, with the peculiarity that they are very long and move in a whip-like fashion, as occurs in sperm cells [11]. Motile cilia (and flagella) contain a 9+2 axoneme consisting of 9 concentric MT doublets and a central MT pair. Along the doublets reside the axonemal dynein molecules whose sliding along adjacent MTs powers the motion of these organelles [3].

Unlike motile cilia and flagella, primary cilia mainly function as cell type-specific signaling organelles. They typically contain a 9+0 axoneme, lacking not only the central MT pair but also axonemal dynein, rendering these cilia immotile. Despite these differences, primary cilia are structurally similar to motile cilia, also emanating from the centrosome’s membrane-docked mother centriole (known as basal body in the ciliary context), and sharing many of the same markers, like acetylated tubulin (AcTub) and ARL13B [12-14]. Among the many important functions of primary cilia are: (i) embryonic Hedgehog signaling to pattern the nervous system and skeleton; (ii) monitoring of blood and urine flow; (iii) sensory perception of light, smell and sound; and (iv) neuropeptide signaling in the brain to control behaviors such as feeding [15-18]. In accordance with their many important functions, motile and/or primary cilia malfunction is associated with genetic diseases called ciliopathies, which can be severe and affect multiple organs such as eyes, kidneys, brain, lungs or skeleton. Besides ciliopathies, primary cilia are also involved in cancer, with constitutive activation of ciliary Hedgehog signaling being a common cause of tumorigenesis [19, 20].

Cilium assembly (ciliogenesis) and disassembly are dynamically regulated with respect to cell cycle progression [21]. In G1/G0 somatic cells, the mother centriole presents subdistal and distal appendages, whereas the daughter centriole does not [22, 23]. While subdistal appendages are involved in MT anchoring, distal appendages contain proteins, like CEP164 (Centrosomal protein of 164 kDa), that are essential for membrane docking of the mother centriole, thus forming the basal body and allowing subsequent ciliogenesis [22, 24, 25]. While primary cilia are typically stable in terminally differentiated cells, ciliated cycling cells must disassemble their cilia in G2, thereby releasing their centrioles from the ciliary base and allowing them to function as MTOCs at the spindle poles during cell division [26]. This is the case, for instance, of several kinds of stem and progenitor cells, whose primary cilia regularly assemble and disassemble in each cell cycle [27].

While the organization and function of cilia have been studied in many somatic cell types, little is known about its role, and even its presence, in germ cells. Spermatogenesis is the process by which males produce their gametes. It comprises at least three functional processes: i) proliferation of spermatogonia (stem germ cells), ii) meiosis of spermatocytes, and iii) maturation of spermatids, which will be transformed into mature spermatozoa (spermiogenesis) [28]. Both meiosis and spermiogenesis require a tight regulation of the cytoskeleton structures, particularly the centrosomes. Meiosis is a specialized cell division that generates haploid gametes from diploid spermatogonia. It comprises two consecutive cell divisions following a single round of DNA replication [29]. To allow these two meiotic consecutive divisions, each one requires the assembly of a spindle, and therefore centrosomes must duplicate twice [30]. After meiosis, spermatids undergo dramatic morphological changes in order to be transformed into fully competent motile spermatozoa [31]. During this process, the sperm tail grows out from the mother centriole of the spermatid, which is thereby transformed into the basal body of the flagellum [32, 33].

Ciliopathies are often associated with infertility [2, 34]. Moreover, some studies have revealed that dysfunction of primary cilia is associated with impaired male reproductive tract development. However, these effects were attributed to the malfunction of the primary cilium in somatic cells of the testicle, like Sertoli and peritubular myoid cells, or cells of the epididymis [35]. More recently, the presence of cilia in meiotic cells has been reported in zebrafish oogenesis [36] and spermatogenesis [37]. These reports indicate that the meiotic cilium may be an important regulator of chromosome polarization and recombination. In addition, these recent findings showed that cilia are present in male and female meiosis in mouse [36], but a morphological or functional description is lacking.

In this study we present the first thorough description of cilia in male mouse meiosis. We uncovered the precise stage at which the cilium is assembled, revealing that meiotic cilia are very transient structures. In addition, and contrary to zebrafish, our results suggest that the role of the cilium in mammalian meiosis is not related to chromosome ends polarization, but instead may be linked to spermatocyte cyst organization.

## RESULTS

### Mouse meiotic cilia are present in zygotene spermatocytes prior to centrosome duplication

The presence of ciliary structures in mouse seminiferous tubules has been recently reported [36]. However, the precise stages at which cilia develop remain unclear. We here present a detailed study of the localization and dynamics of the mouse meiotic cilium. For this purpose, we first studied the localization of the main component of its axoneme, acetylated tubulin (AcTub), during both meiotic divisions in squashed spermatocytes.

In early prophase I, at leptotene, AcTub labels the centrioles colocalizing with Centrin 3 (CETN3) (Figure 1A and A’). In the transition from leptotene to zygotene, two alternative patterns could be seen. Some spermatocytes showed elongated hair-like structure labelled with AcTub, emanating from one of the CETN3 centriole signals (Figure 1B and B’), while some others presented AcTub only at the centrioles (Figure 1C and C’). Quantitative analysis showed that around 20% of the spermatocytes at zygotene showed this hair-like structure (n=100 spermatocytes at zygotene per individual, three biological replicates). This hair-like staining pattern of AcTub in spermatocytes at early zygotene is consistent with the solitary projection that is formed when primary cilia assemble in both somatic [12] and meiotic cells [36]. These zygotene cilia were present when only two centrioles are detected, which indicates that mouse meiotic cilia are assembled prior to centrosome duplication.

**Figure 1.**
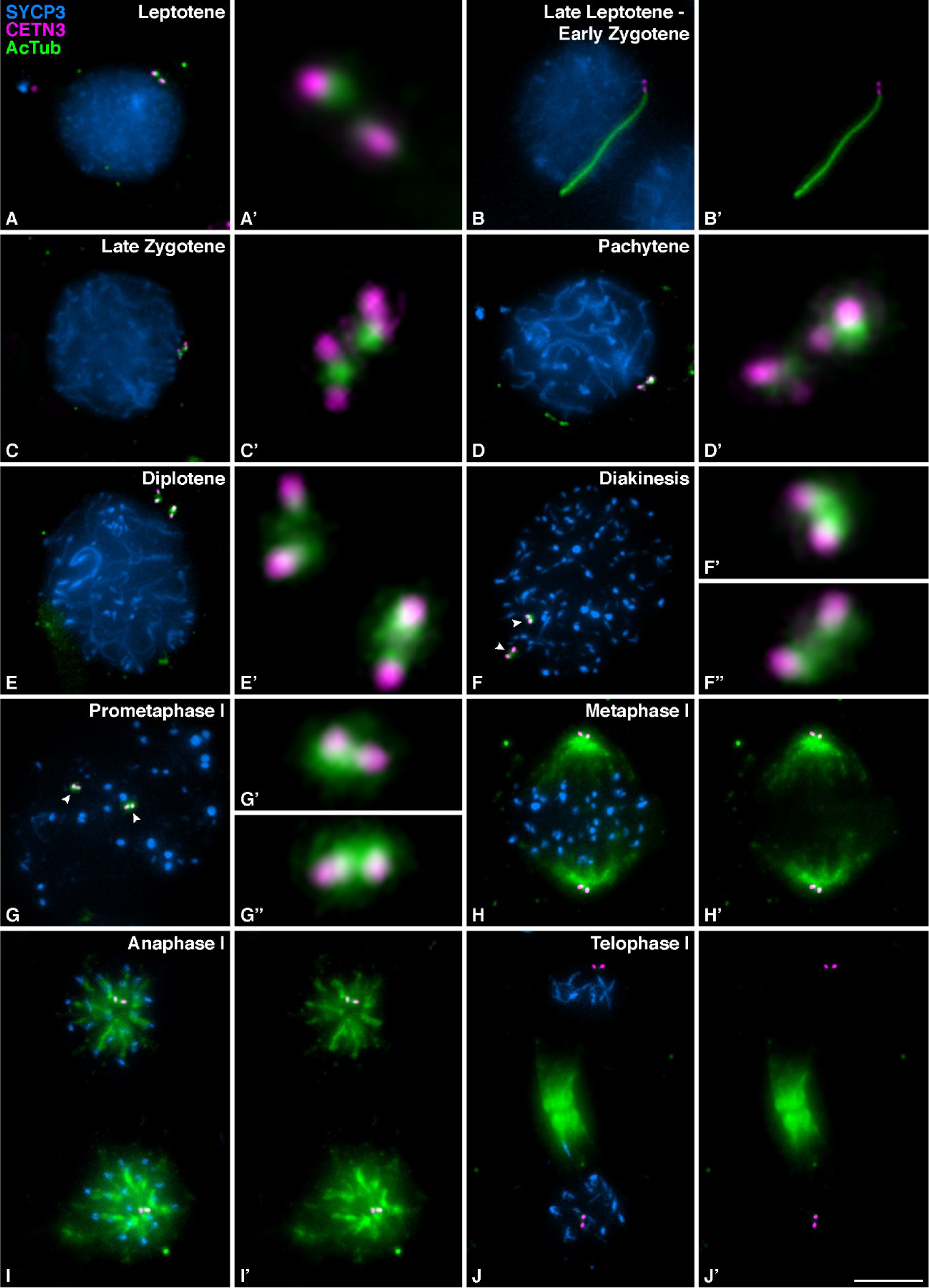
Mouse spermatocytes form transient cilia during meiotic prophase I. Detection of acetylated Tubulin in mouse spermatocytes during the second meiotic division. Triple immunolabelling of SYCP3 (magenta), Centrin 3 (CETN3) (yellow) and acetylated Tubulin (AcTub) (green) on squashed WT mouse spermatocytes at (A) Leptotene, (B) Leptotene to zygotene transition, Late leptotene - Early Zygotene, with cilium, (C) Late Zygotene, (D) Pachytene (E) Diplotene, (F) Diakinesis, (G) Prometaphase I, (H) Metaphase I, (I) Anaphase I, (J) Telophase I. For images A’ and C’-G’ the 300X magnification for centrosomes is shown (white arrows). Scale bar in J’ represents 5 μm.

Centrioles duplicate during zygotene [30], as revealed by the presence of four different signals labeled with CETN3. At pachytene, AcTub is detected as a single signal per centrosome, associated to the duplicated centrioles (Figure 1D and D’). At diplotene (Figure 1E and E’) and diakinesis (Figure 1F and F’-F’’), when centrosomes start migrating towards opposite poles, the signal of AcTub expands to surround the two pairs of separated centrioles. During prometaphase I, AcTub still localizes to the centrosomes (Figure 1G and G’-G’’). Once centrosomes have completely migrated to opposite poles and the meiotic bipolar spindle is established at metaphase I, AcTub labels the two pairs of centrioles and also the MTs of the bipolar spindle, while CETN3 only detects the centrioles (Figure 1H and H’). The same pattern persists at anaphase I, when homologous chromosomes segregate (Figure 1I and I’). At telophase I, CETN3 still labels the centrosomes and AcTub labels the centrioles and the midzone MTs (Figure 1J and J’). During interkinesis, AcTub signal again accumulates at the centrosomes, as duplicated centrioles are clearly seen by the four CETN3 signals upon the second centrosome duplication (Figure 2A and A’). At metaphase II (Figure 2B and B’) and anaphase II (Figure 2C and C’), AcTub is observed at the centrioles and the meiotic bipolar spindle II. At telophase II, AcTub labels the centrioles and the midzone MTs (Figure 2D and D’).

**Figure 2.**
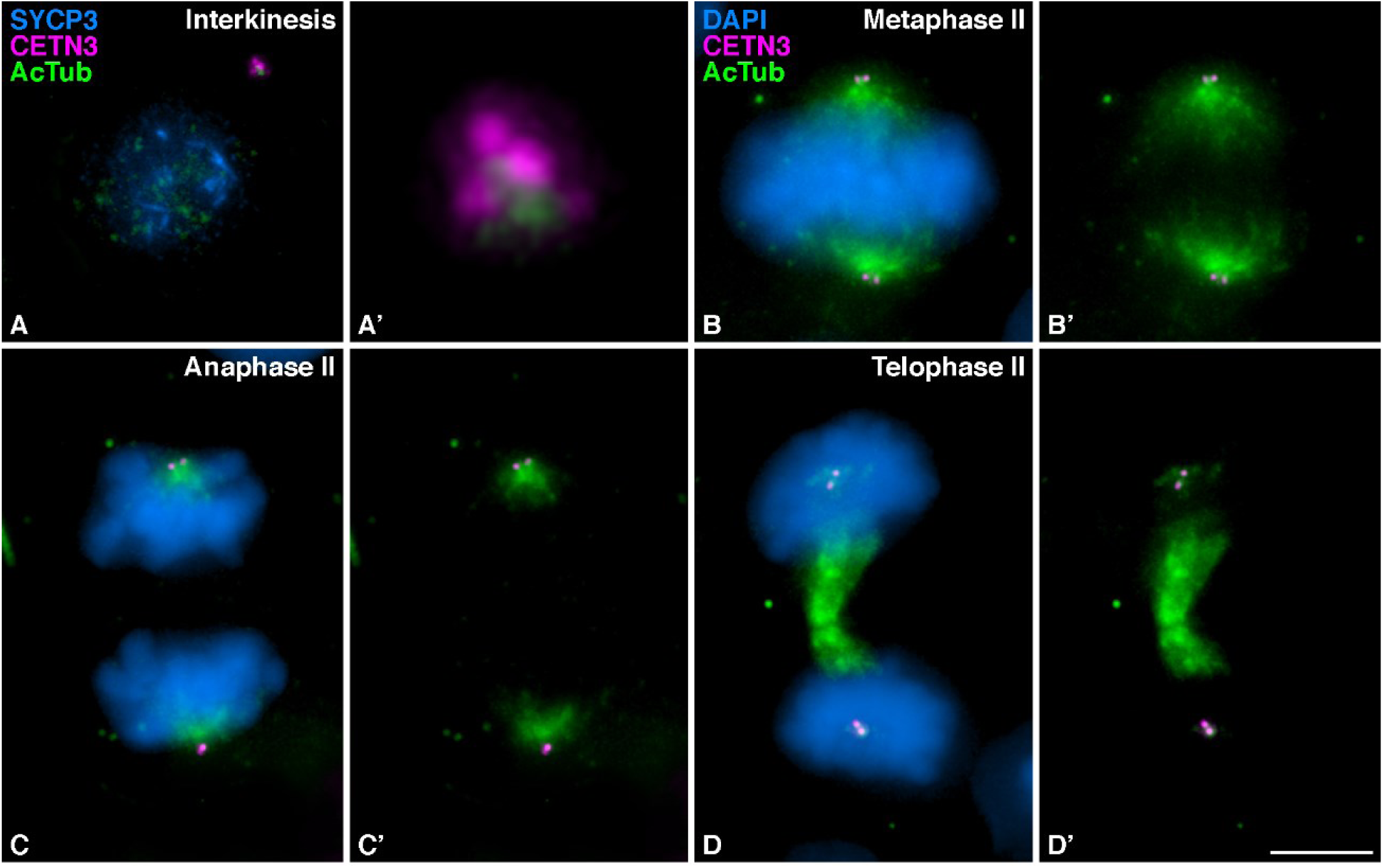
Mouse secondary spermatocytes do not present cilia. Detection of acetylated Tubulin in mouse spermatocytes during the second meiotic division. Triple immunolabelling of SYCP3 (magenta), Centrin 3 (CETN3) (yellow) and acetylated Tubulin (AcTub (green) on squashed WT mouse spermatocytes at (A) Interkinesis. And double immunolabelling of Centrin 3 (CETN3) (yellow) and acetylated Tubulin (AcTub (green), with chromatin stained with DAPI (blue) at (B) Metaphase II, (C) Anaphase II (D) Telophase II. For images A’ the 300X magnification of the centrosomes is shown. Scale bar in D’ represents 5 μm.

Tubulin is a protein whose distribution has been extensively studied in mammalian meiosis. Therefore, it was surprising that the meiotic cilium was not observed before. In order to compare the distribution of AcTub with that of overall tubulin, we performed a co-labeling with an antibody against αTubulin (αTub). A careful comparison of non-acetylated αTub and AcTub lead us to conclude that both proteins show a different localization pattern in meiosis (Figure 3). At leptotene, αTub is mainly localized to cytoplasmatic MTs, while AcTub accumulates only at the centrosome (Figure 3A). In those zygotene spermatocytes in which the cilium was present, AcTub labeled this structure, while αTub is only located along cytoplasmatic MTs (Figure 3B). This indicates that αTub does not label meiotic cilia. This may explain why these structures were not detected in previous studies. At metaphase I and metaphase II, both non-acetylated and acetylated tubulin labelled the spindle MTs, but only AcTub is detected at the centrosomes (Figure 3C,D).

**Figure 3.**
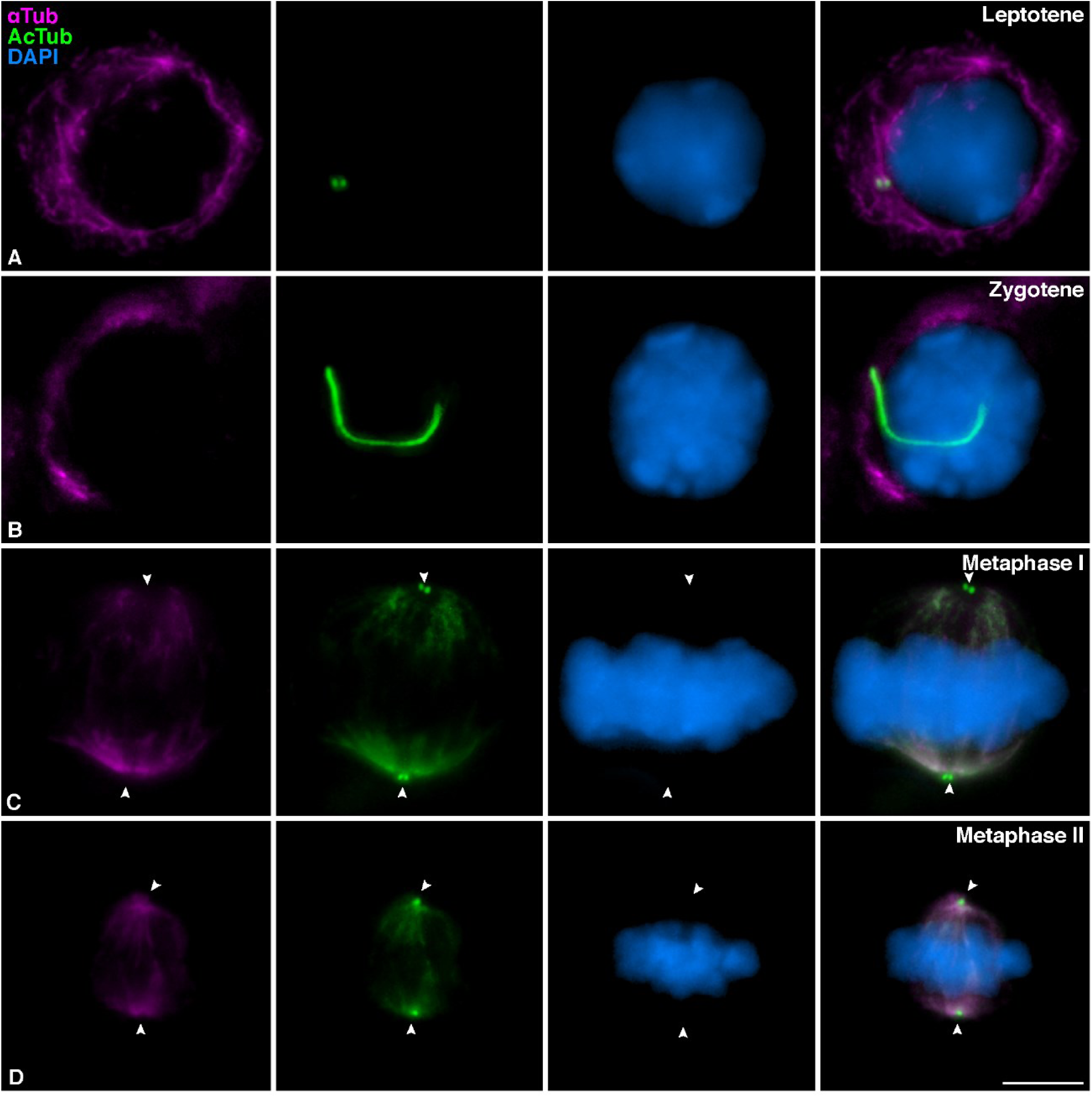
Mouse spermatocyte cilia are detected with AcTub but not alpha-Tub antibodies. Double immunolabelling of α Tubulin (αTub) (green) and acetylated Tubulin (AcTub) (magenta), with chromatin stained with DAPI (blue) on squashed WT mouse spermatocytes at (A) Leptotene, (B) Zygotene with primary cilium, (C) Metaphase I, (D) Metaphase II. White arrows in C and D indicate centrosomes localization. Scale bar in D represents 5 μm.

To further characterize the organization and composition of the mouse meiotic cilium we studied the localization of AcTub and ADP-ribosylation factor-like 13B (ARL13B) in histological cryosections of mouse testis. ARL proteins belong to the ARF family of GTPases, which are involved in ciliogenesis and ciliary intraflagellar transport (IFT) in somatic cells [38, 39]. More specifically, ARL13B mediates primary cilia function and/or Hedgehog signaling [40]. In testis sections, spermatocytes located at the base of the seminiferous tubule were the only ones that presented cilia and SYCP3 signal compatible with early prophase I (Figure 4 I.Aa-d). However, AcTub also labeled flagella of spermatids facing the lumen of the seminiferous tubule (Figure 4 I.A). ARL13B was present along the meiotic cilia visualized at prophase I spermatocytes located at the base of the tubule (Figure 4 I.Ba-d). Triple immunolabeling of ARL13B, AcTub and SYCP3 in squashed spermatocytes corroborated that ARL13B and AcTub colocalized in zygotene spermatocytes (Figure 4 II.A-D).

**Figure 4.**
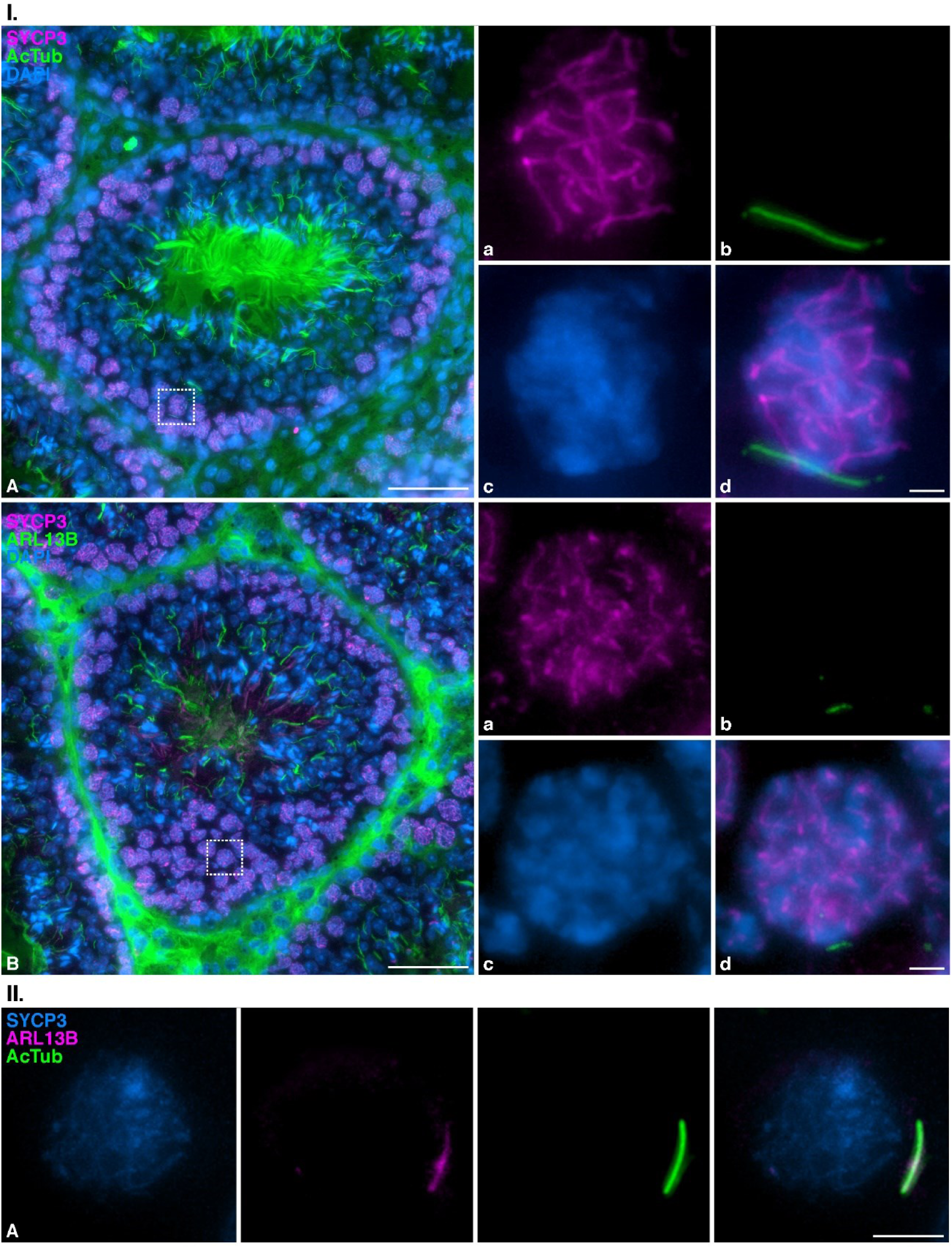
Zygotene cilia of mouse testis spermatocytes contain ARL13B. **I. Distribution of acetylated Tubulin and ARL13B in mouse testis cryosections** Double immunolabelling of SYCP3 (magenta) and acetylated Tubulin (AcTub) (green), with chromatin stained with DAPI (blue) on cryosections of mouse testis. (A) complete section of a seminiferous tubule, (a-d) amplified selected spermatocyte at zygotene showing a polymerized primary cilium. And double immunolabelling of SYCP3 (magenta) and ARL13B (green), with chromatin stained with DAPI (blue) on cryosections of mouse testis. (B) complete section of a seminiferous tubule, (a-d) amplified selected spermatocyte at zygotene showing a incipient polymerizing primary cilium. Scale bar in A and B represents 50 μm and scale bar in d represents 5 μm. **II. Distribution of ARL13B in mouse squashed spermatocytes at zygotene** Triple immunolabelling of SYCP3 (magenta), ARL13B (green) and acetylated Tubulin (AcTub) (yellow) on squashed WT mouse spermatocytes at zygotene (A). Scale bar represents 5 μm.

These results confirm that the fiber-like structures that we observe at zygotene spermatocytes are *bona fide* cilia. Nevertheless, given that mouse flagella also incorporate AcTub [11], an additional corroboration was conducted in order to discard that the cilia observed in zygotene could be mistaken with the flagella of nearby spermatozoa after the squashing procedure. For this, we studied the localization of AcTub in spermatocytes of *Ankrd31*^*–/–*^ [41], a sterile mouse model that lacks spermatozoa. Our results showed that *Ankrd31*^*–/–*^ present zygotene cells with AcTub-labelled cilia (Figure A1 A,B), with a similar morphology than wild type (WT) individuals (Figure A1 C,D). In contrast, only WT mice present early spermatids (Figure A1E) and elongated mature spermatids (Figure A1 F) with AcTub-labelled flagella.

In conclusion, all these results lead us to conclude that cilia are present in mouse spermatocytes at zygotene prior to centrosome duplication.

### Meiotic cilia appear at the onset of synapsis

We next wanted to assess the temporal correlation between the formation of this structure and other typical meiotic processes, such as chromosome synapsis. For this, we triple immunolocalized AcTub with SYCP3 and SYCP1, a component of the transverse filaments of the synaptonemal complex [42]. At leptotene, when synapsis has not yet started and SYCP1 labelling is absent, AcTub is only detected at the centrioles (Figure 5A). The cilium is first observed in late leptotene spermatocytes, when SYCP3-labeled axial elements are already formed but SYCP1 protein is not yet loaded to chromosomes. The length of the cilium at this stage is approximately 5 μm, presumably polymerizing (Figure 5B). Synapsis initiation marks the beginning of zygotene. At this stage, short filaments of SYCP1 were seen partially colocalizing with SYCP3. The cilium elongated conspicuously at this stage, reaching a maximum length of 15μm (Figure 5C). When synapsis has partially progressed and longer filaments of SYCP1 are clearly seen, the meiotic cilium seems to start shrinking, being around 10-13 μm long at mid zygotene (Figure 5D) and late zygotene (Figure 5E). As previously indicated, the cilia were present in only a fraction (≈ 20%) of zygotene cells. Therefore, most spermatocytes at this stage did not present cilia. In those cases, AcTub only labelled centrioles (Figure 5F). These results led us to conclude that mouse meiotic cilia start to polymerize at the transition from leptotene to zygotene, and they are fully polymerized by mid zygotene.

**Figure 5.**
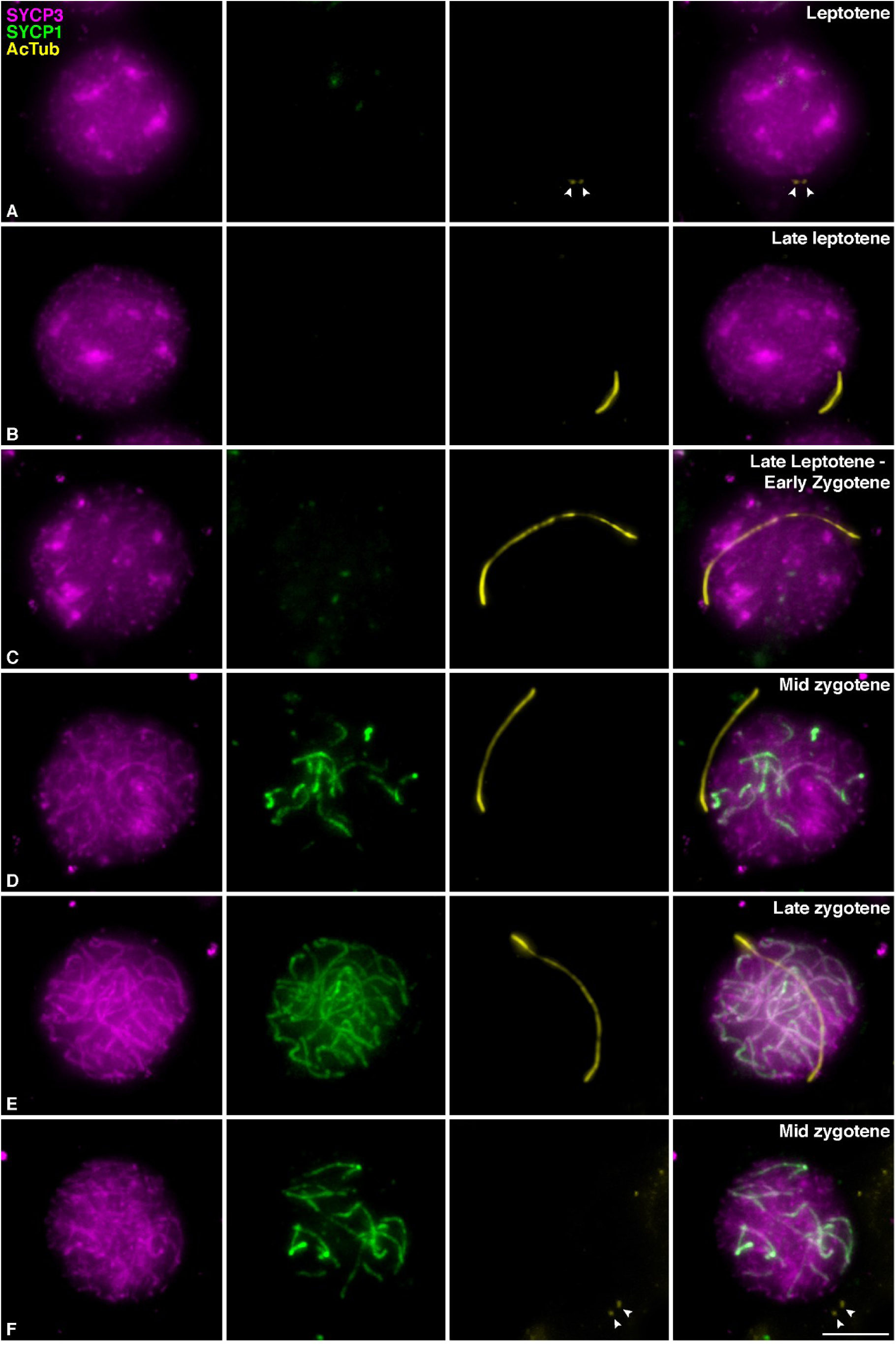
Mouse meiotic cilia appear at the onset of synapsis. Triple immunolabelling of SYCP3 (magenta), SYCP1 (green) and acetylated Tubulin (AcTub) (yellow) on squashed WT mouse spermatocytes at (A) Leptotene, (B) Late Leptotene with polymerizing primary cilium, (C) Leptotene to zygotene transition, Late leptotene - Early Zygotene with primary cilium, (D) Mid Zygotene with primary cilium, (E) Late Zygotene with primary cilium, (F) Mid zygotene without cilium. Scale bar in F represents 5 μm.

### Mouse meiotic cilia emanate from the mother centriole prior to centrosome duplication

We then tested if mouse meiotic cilia emanate from the mother centriole. For this purpose, we first studied the dynamics of CEP164, a protein of the distal appendages required for cilia formation in somatic cells [22, 43] that has also been identified at the mother meiotic centriole in mouse spermatocytes [44]. Our results corroborated that CEP164 labeled one of the two centrioles before centrosome duplication at early prophase I. At leptotene, CEP164 labels a circular signal around one of the two CETN3-labeled centrioles of the not yet duplicated centrosome (Figure 6 I. A and A’). Centrosome duplication occurs during zygotene [30, 44], and four signals of CETN3 were observed, but CEP164 still labeled only one of the four centrioles at this stage (Figure 6 I.B and B’). We then we triple immunolabelled CETN3, CEP164 and AcTub. We found that the cilia emanated from the centriole that possessed CEP164 at early zygotene, i.e, from the mother centriole of the not yet duplicated centrosome (Figure 6 II.A). CETN3 signal did not overlap with AcTub cilium labeling, suggesting that when the cilium is polymerized, the centrioles are not acetylated (Figure 6 II.A’). These results indicate that mouse meiotic cilia appear at late leptotene-early zygotene transition when centriole duplication has not yet occurred, and that cilium emanates from the mother centriole.

**Figure 6.**
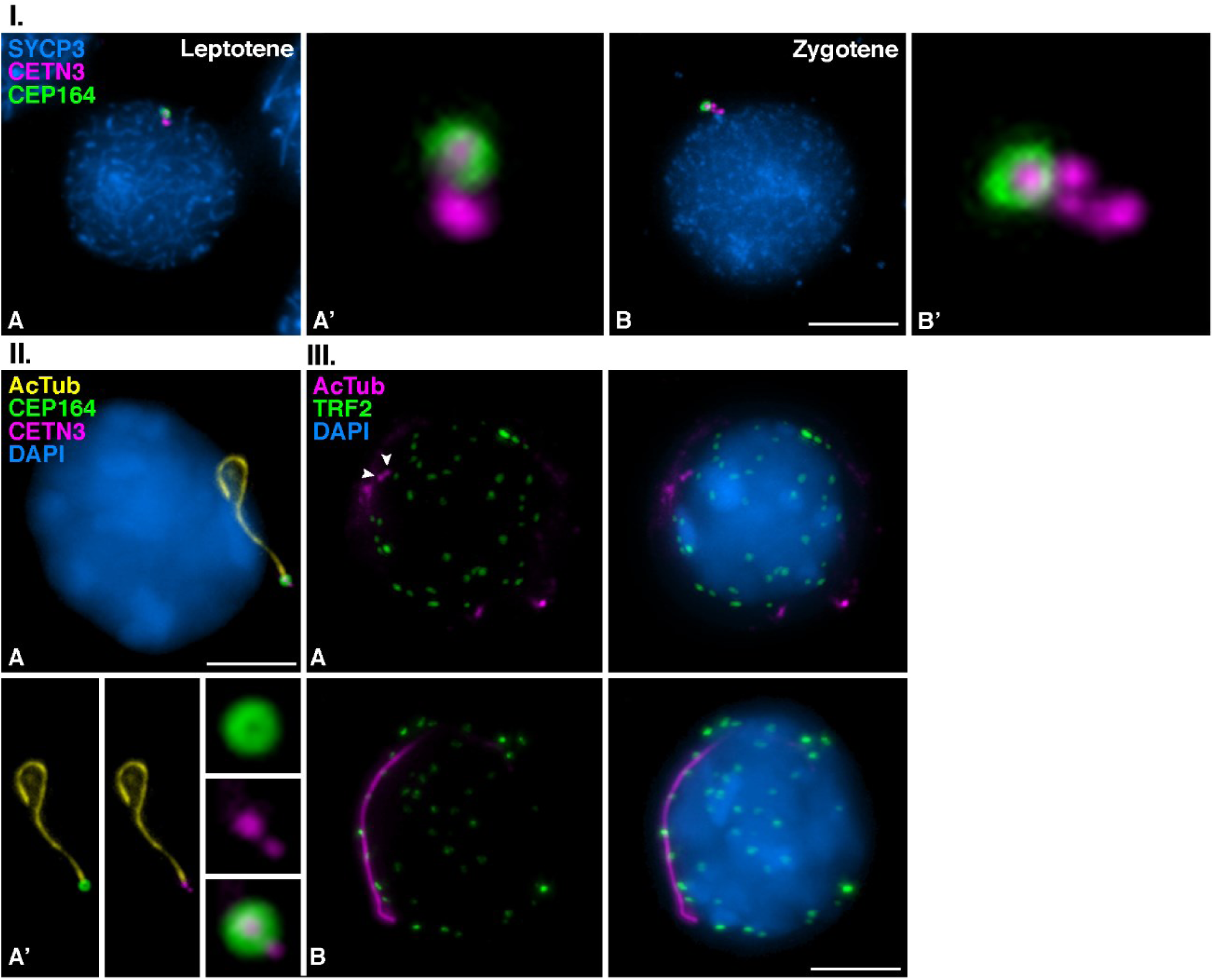
Mouse meiotic cilia emanate from the mother centriole before centrosome duplication. **I. Distribution of CEP164 in early prophase I spermatocytes** Triple immunolabelling of SYCP3 (blue), Centrin 3 (CETN3) (red) and CEP164 (green) in mouse spermatocytes at (A) Leptotene, (B) Zygotene. Images A’-B’show the 300X magnification of the centrosomes. Scale bar in H represents 5 μm. **II. Meiotic zygotene cilia emanates from the unduplicated mother centriole** Triple immunolabelling of acetylated Tubulin (AcTub) (yellow), CEP164 (green) and Centrin 3 (CETN3) (magenta), with chromatin stained with DAPI (blue) on squashed WT mouse spermatocytes at (A) Zygotene presenting a primary cilium. Images in A’ show the primary cilia and the 300X magnification of the centrosomes. Scale bar in A represents 5 μm. **III. The presence of the primary cilia is not directly related to the *bouquet* conformation of chromosome ends** Double immunolabelling of acetylated Tubulin (AcTub) (magenta) and TRF2 (green), with chromatin stained with DAPI (blue) on squashed WT mouse spermatocytes at (A) Zygotene without primary cilium, (B) Zygotene presenting a primary cilium. Scale bar in B represents 5 μm.

In order to corroborate the pattern of distribution of CEP64 and its implication during male mouse meiosis, we continued studying its localization during both meiotic divisions. We observed that CEP164 is sequentially loaded to the centrioles until the four centrioles are assembling distal appendages at the end of the first meiotic division (Figure A2 A-F), according to previous results spermatocytes [44]. There is no incorporation of additional distal appendages in the daughter centrioles during the second meiotic division. CEP164 labeled only one of the two centrioles per centrosome in opposite poles of the bipolar meiotic spindle II. When spermiogenesis progressed, centrioles were disorganized in late spermatids, no CETN3 signals were seen, however, the CEP164 signal is still present as previously reported [44, 45], indicating the formation of a basal body for the growing flagellum (Figure A2 A-E).

### Meiotic *bouquet* conformation and cilia formation are not concurrent events

Meiotic cilium in zebrafish oocytes favors the polarization of the chromosome telomeres, which are anchored to the nuclear envelope (NE), forming the so-called *bouquet* configuration [36]. To test whether the mouse zygotene cilium in spermatocytes is related to the *bouquet* conformation, we performed a double immunodetection of the telomeric double-stranded TTAGGG repeat binding proteins (TRF2) and AcTub. In spermatocytes at zygotene, we observed TRF2 as small and rounded signals randomly distributed, presumably over the NE, regardless of whether cilia labelled by AcTub is present (Figure 6 III.A) or absent (Figure 6 III.B). Therefore, there seems not to be a direct relationship between the presence of a cilium and the configuration of the telomeric *bouquet* in mouse spermatocytes.

### Zygotene TEX14 connected cysts

Classical studies on the organization of the seminiferous tubules showed that spermatocytes form cysts of cells synchronously advancing through meiosis. These spermatocytes present cytoplasmic bridges that presumably provide a biochemical and physiological coordination of the cells in a cyst. As our results showed that only ≈ 20% of the spermatocytes at zygotene have a polymerized cilium, we then wanted to test if spermatocytes displaying the cilium could form specific cysts. For this, we detected TEX14, the product of testis-expressed gene 14 (*Tex14*), a protein that localizes to germ cell intercellular bridges in male spermatocytes [46]. Our results showed that TEX14 labeled ring-shaped intercellular bridges between spermatocytes at different stages. Cysts of early prophase I spermatocytes are seen at the base of the seminiferous tubules in testis cryosections (Figure 7 I.Aa). In most of these spermatocyte cysts, the cilium could be detected in only one of the spermatocytes (Figure 7 I. Ab).

**Figure 7.**
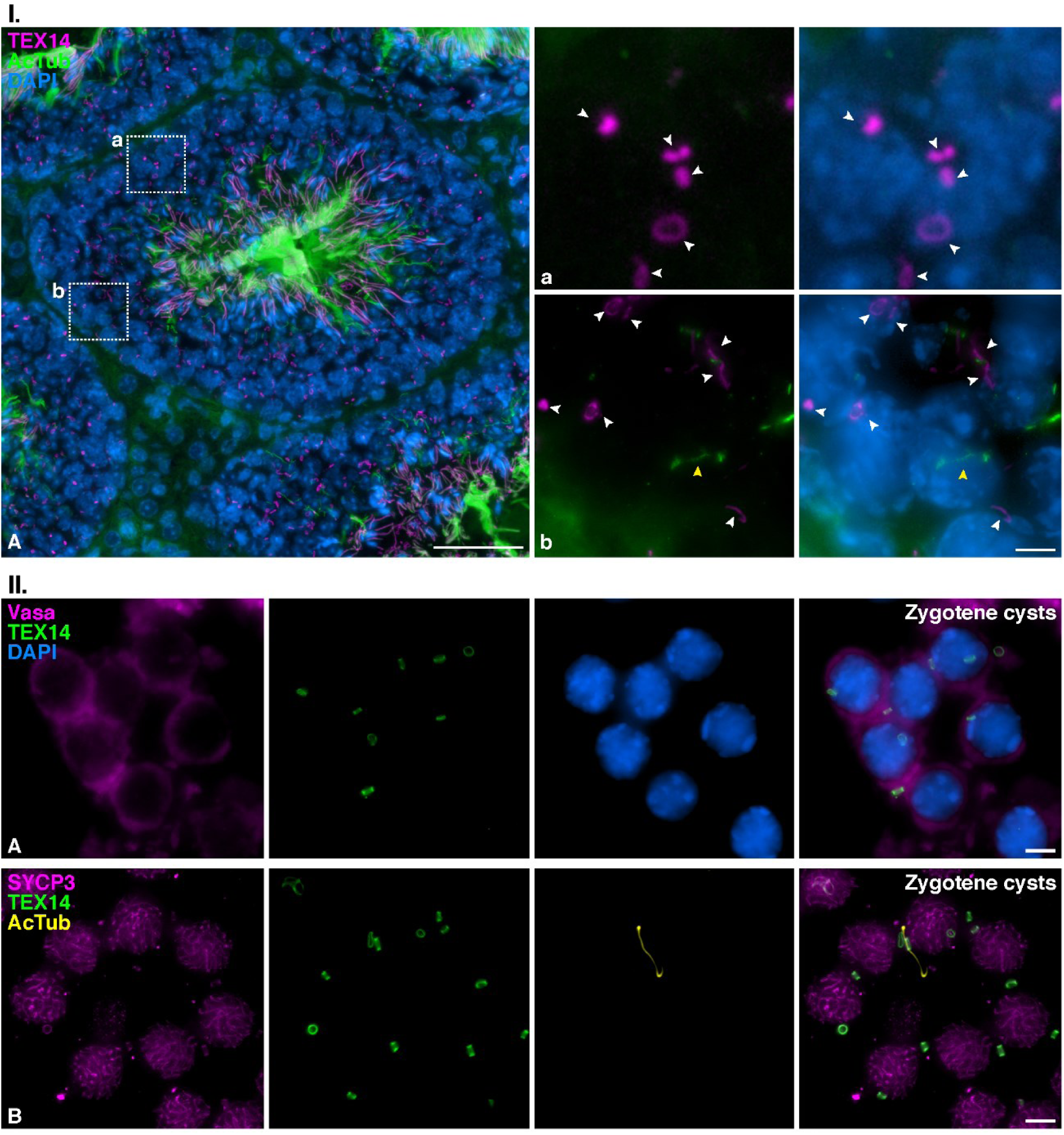
Spermatocyte cysts share cilia among several spermatocytes at zygotene. **I. Distribution of TEX14 cellular bridges in mouse testis cryosections**. Double immunolabelling of TEX14 (magenta) and acetylated Tubulin (AcTub) (green), with chromatin stained with DAPI (blue) on cryosections of mouse testis. (A) complete section of a seminiferous tubule. Scale bar in A represents 50 μm. (a) amplified cyst of primary spermatocytes interconnected by TEXT14 bridges (white arrows), (b) amplified cyst of primary spermatocytes interconnected by TEXT14 bridges (white arrows), with one zygotene showing a primary cilium (yellow arrow). Scale bar in d represents 5 μm. **II. Distribution of TEX14 cellular bridges in mouse spermatocytes at zygotene** (A) Double immunolabelling of VASA (magenta) and TEX14 (green), with chromatin stained with DAPI (blue) on squashed WT mouse spermatocytes. A cyst of spermatocytes at zygotene interconnected by TEXT14 bridges is shown. Scale bar represents 5 μm. (B) Triple immunolabelling of SYCP3 (magenta), TEX14 (green) and acetylated Tubulin (AcTub) (yellow). A cyst of spermatocytes at zygotene interconnected by TEXT14 bridges is shown, with one zygotene showing a primary cilium. Scale bar represents 5 μm.

The association of spermatocytes in cysts was also preserved in preparations of squashed seminiferous tubules. We double immunolabeled TEX14 and VASA, a germ specific cytoplasmic marker [47] and corroborated that there is a cytoplasmic continuity between the spermatocytes at the places where TEX14 bridges are present (Figure 7 II. A). Our results showed that not all the cells within the same cyst presented cilia (Figure 7 II. B). This demonstrates that cells bearing a cilium are not specifically connected to each other in a cyst, but on the contrary, they preferentially form connections with other spermatocytes lacking the cilium.

## DISCUSSION

### The meiotic cilium in mouse spermatogenesis

The primary cilium is an outgrowth of the plasma membrane located in many eukaryotic cell types. These immobile and solitary projections have very diverse functions related to cell communication [48], and have traditionally been depicted as structures whose continued presence is incompatible with cell cycle progression [26]. However, although studies on the structure of primary cilia in various somatic tissues are extensive and diverse, their presence in vertebrate meiosis has only been reported in two recent works. The first one described the zygotene cilium in zebrafish oocytes, pointing to an evolutionary conservation of this structure in mammals, but suggesting that ciliary structures in mouse spermatocytes were very short, barely extending beyond the basal body [36]. The second study proposed that germ cell-specific depletion of ciliary genes resulted in compromised double-strand break repair, reduced crossover formation, and increased germ cell apoptosis in zebrafish spermatogenesis [37]. We here show a detailed description of the mouse meiotic cilium, and we hypothesize that they are primary cilia. We found that around 20% of the spermatocytes at zygotene present cilia as a solitary long protrusion that disassembles once spermatocytes progress into pachytene. This provides a scenario that challenges the assumption that primary cilia are antagonistic with cell division, as it has to be taken into account that cells at zygotene have already started meiotic cell division [49]. Apart from our work and the recent advances in zebrafish gametogenesis [36, 37], another previous report also pointed at the presence of primary cilia during *D. melanogaster* spermatogenesis. Moreover, they suggested that these cilia were stable through both meiotic divisions in dividing spermatocytes with yet unknown functions [50]. Therefore, it seems that, although somatic primary cilia in vertebrates are a sensory machinery during interphase which was considered a “sleeping beauty” to date during cell division [51, 52], cells that undergo meiosis are capable of polymerizing cilia once cell division has started. This may be related to the dynamics of the mitotic versus the meiotic centrosome. In somatic cells, centrosome duplication occurs, like DNA duplication, during S phase, i.e. before mitosis [53]. In contrast, during gametogenesis, DNA duplication occurs in the premeiotic interphase, whereas centrosome duplication happens twice once meiosis has begun: in mouse, at zygotene and again at interkinesis [30, 44]. This implies that somatic centrosome duplication occurs after ciliogenesis and before entry into mitosis, whereas meiotic centrosome duplication occurs after ciliogenesis, with both processes taking place in the course of meiosis.

### Meiotic centrosomes and their relation to ciliogenesis and flagelogenesis

Our results demonstrate that zygotene cilia in mouse spermatocytes contain acetylated Tubulin -AcTub-, similarly to the zygotene cilia in zebrafish spermatocytes and oocytes [36, 37]. This was expected since this posttranslational modification of Tubulin is a well-known marker in both motile and primary cilia in somatic cells [54-56]. We detected that tubulin is also acetylated in the flagellar axoneme of spermatids, as expected [11]. Given that the cilium emanates from the larger rounded signal of the two centrioles marked by CETN3, where CEP164 distal appendages are assembled, we conclude that zygotene cilia emanate from the mother centriole, as in somatic cells [57, 58]. CEP164 distribution is consistent with the function of distal appendages in cilia and flagella [59, 60]. On the one hand, we show that CEP164 serves to dock the basal body during zygotene ciliogenesis, sharing a similar role to their function in somatic cells [22, 24, 25]. On the other hand, spermatids present CEP164 distal appendages to promote the polymerization of the flagellar axoneme, as previously reported [44]. Hence, the dynamics of CEP164 seems to be essential for the progression of spermatogenesis. This is consistent with the sterile phenotype of CEP164 deficient mice -FoxJ1-Cre;CEP164fl/fl-that showed sperm agglutination and aberrant ciliogenesis of motile multicilia in the somatic cells of the efferent ducts [45]. With the present work, we now suggest that CEP164 not only participates in flagelogenesis, but also in mouse meiotic ciliogenesis. For further clarification, we present a scheme representative of the progression of male mouse meiosis in relation to centrosome dynamics, ciliogenesis and flagelogenesis (Figure 8).

**Figure 8.**
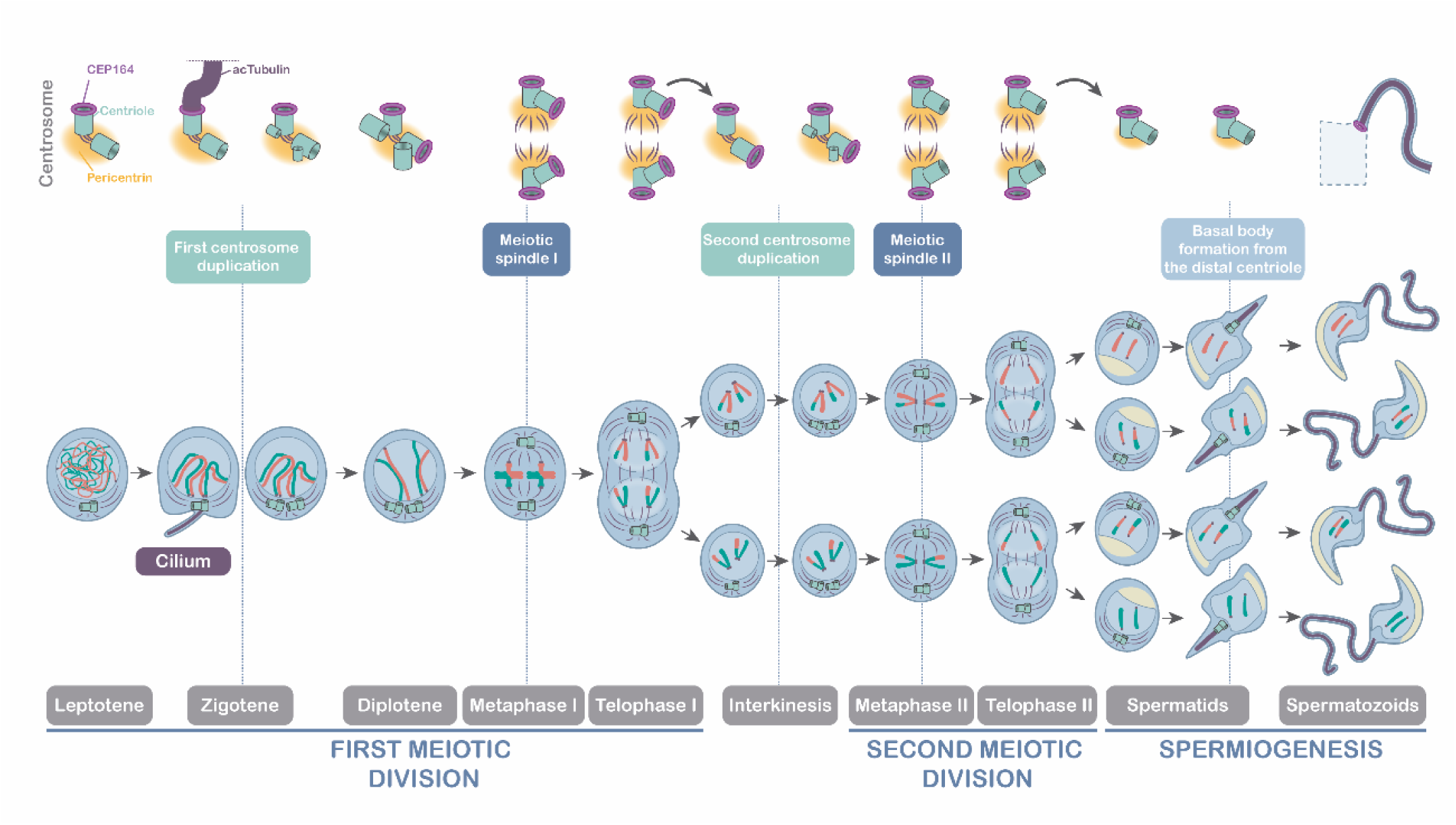
Schematic representation of the meiotic stages in relation to centrosome dynamics. Meiosis progression showing the centrosomal events (centrosome duplication I and II and formation of meiotic spindle I and II). Ciliogenesis and flagelogenesis events are represented: Centrosome duplication occurs at zygotene transition and interkinesis; The formation of primary cilia occurs at zygotene transition; Flagellum is formed in spermatids. The distribution of centrioles, pericentriolar matrix (Pericentrin) and distal appendages (CEP164) is represented thorough both meiotic divisions. Primary cilia at zygotene and flagella at spermatids is represented with acetylated Tubulin (AcTub).

### Tubulin acetylation during male mouse meiosis

We here showed that centrosomes present AcTub partially colocalizing with CETN3 during the entire meiotic division, but non-acetylated α-Tub is never detected at centrosomes. CETN3 is a component of meiotic centrioles [30, 44], therefore we can conclude that meiotic centrioles possess acetylated tubulin. This might promote their stability, since experiments using biotinylated tubulin demonstrated that centriolar MTs do not exhibit significant exchange with the cytoplasmic protein pool [61]. This agrees with centriolar MTs being heavily acetylated and polyglutamylated, which are hallmarks of stable MTs [62]. In this sense, acetylation of MTs promotes the recruitment of PLK1 to the centrosomes in mitosis [63]. This kinase is located at meiotic centrosomes regulating their migration [30, 44]. Interestingly, our results also conclude that when meiotic cilia are polymerized, centrioles do not show AcTub. Consequently, future research should clarify the relationship between PLK1 and the acetylation of centriolar tubulin, and the role that this kinase might exert on meiotic ciliogenesis.

On the other hand, during mitotic metaphase, AcTub is enriched at interpolar and kinetochoric MTs, but not at astral MTs, and AcTub becomes concentrated on the midbody during telophase and cytokinesis [64]. This acetylation is an evolutionarily conserved posttranslational modification of α-Tub associated with increased stability of various MT populations, including the mitotic spindle [56]. Moreover, tubulin acetylation does modulate the ability of mitotic MTs to bind to MAPs (Microtubule Associated Proteins) and motor proteins, and may regulate MT stability and function [65, 66]. Our results show that once the bipolar spindles are shaped at metaphase I and II, spindle MTs are acetylated. We also observed labelling of AcTub in the midbody of telophase I and II. It has been described that the post-translational modifications of tubulin are related to the integrity of the spindle in oocytes [67] and for the stability of the spermatozoa [11]. Specifically, altered acetylation level of tubulin leads to defective spindle assembly and chromosome misalignment in mouse oocyte meiosis [68, 69], but the function of AcTub in male mouse meiosis was not yet described. We suggest that the acetylation of tubulin in meiotic MTs might play similar roles, contributing towards the stability of the centrioles throughout the entire meiosis, and towards the stability of the meiotic spindles I and II at metaphase I/II and the midbodies during telophase I/II.

### Zygotene cilia and *bouquet* formation are not directly related in male mouse meiosis

From our results, it emerges that the cilium appears in primary spermatocytes at the transition from leptotene to zygotene, prior to centriole duplication. This indicates that the cell signaling processes in which the cilium may be involved might be related to the initiation of synapsis [42, 70]. The *bouquet* polarization is a transient conformation of the chromosomal ends that favors the pairing process between homologous chromosomes [71]. Recent reports in zebrafish oogenesis demonstrated a correlation between the zygotene cilia and the *bouquet* conformation, suggesting that absence of cilia or their incorrect formation affect synapsis progression [36]. However, we could not find any correlation between the presence of cilia and the chromosomal *bouquet* conformation in mice. This conformation is extremely short-lived in this species and most likely precedes the formation of the cilia. On the other hand, the zebrafish cilium has also been related to recombination regulation [37]. We cannot make any considerations in this regard, as the relationship between cilia and recombination in mice should require additional approaches.

### Spermatocyte cysts facilitate progression of mouse spermatogenesis

Primary cilia are found in many mouse cell types, and their morphology and size vary depending on their cellular functions. For example, primary cilia of cardiac fibroblasts are essential for heart development [72], and primary cilia of epithelial cells from the renal proximal tubule participate in the regulation of the glomerular filtration barrier [73].

In spermatocytes, we have observed the presence of a remarkably long cilium, approximately 15 μm, compared to the 6 μm described for zebrafish spermatocyte cilia [36]. If being a primary cilium, the length of the cilium in mouse spermatocytes is similar to that of mouse renal epithelial cells, neurons, or olfactory cells [74, 75]. In these somatic cells, primary cilia have a sensory function, capturing information from the extracellular environment and triggering an intracellular response. Thus, greater surface of ciliary membrane could harbor more sensory transmembrane proteins and therefore more signaling receptors. That is why longer ciliary length has been related to greater sensitivity and response [76], and modifications of the ciliary length affect cilium integrity and ultimately its functions [74]. Given that the mouse meiotic cilium has a mean length equivalent to other long cilia in mouse somatic cells, we suggest that it may also perform important cell signaling functions during spermatogenesis. In parallel, we have also shown that mouse zygotene cilia present ARL13B, a GTPase of the ARL family known to mediate ciliary protein trafficking and regulate ciliary Hedgehog signaling [40]. Therefore, it is plausible that mouse zygotene cilia have an active Hedgehog pathway, which should be clarified in detail in future studies.

The histology of the testis might explain the absence of cilia in all the spermatocytes. The epithelia of the seminiferous tubules present cysts of cells that are undergoing spermatogenesis. Spermatogonia are linked by cytoplasmic bridges whose absence leads to infertility [46]. These bridges contain, among other proteins, the testis-expressed gene 14 product (TEX14) [46, 77]. These bridges are partially released once spermatogonia evolve to primary spermatocytes upon entry to meiosis [78]. The proposed roles for the intercellular bridges include germ cell communication and synchronization [79]. The fact that only 20% of the spermatocytes at zygotene present cilia leads us to suggest that zygotene cysts might share a single, or very few, cilia. We have shown that primary spermatocytes are linked by TEX14 rings. Thus, we argue that primary spermatocyte’s cysts do not require cilia in each spermatocyte to progress through spermatogenesis. For further clarification, a scheme representing the seminiferous epithelia is presented (Figure 9).

**Figure 9.**
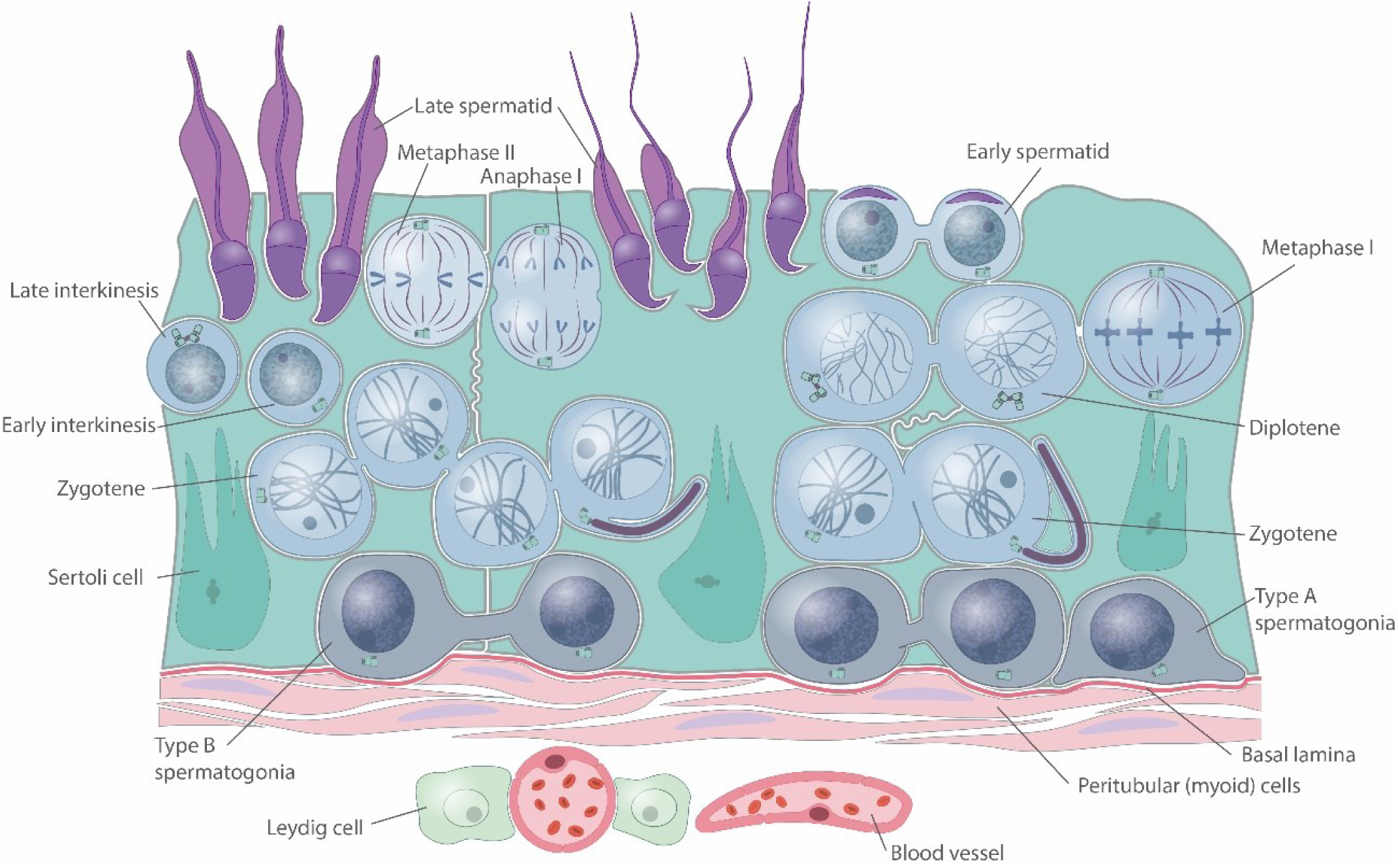
Schematic representation of the seminiferous epithelium. Scheme represents spermatogonias (dark blue), spermatocytes (blue) and spermatids (purple) embedded on the cytoplasm of Sertoli cells (turquoise). Centrosomes are indicated for spermatogonias and spermatocytes. Cilia are represented in one of the zygotenes of a cyst of primary spermatocytes. Peritubular and interstitial compartment is represented indicating the position of myoid cells, Leydig cells and blood vessels.

In conclusion, we can argue that the need to maintain intercellular connections in spermatocyte cysts for achieving fertility [79], together with the unique anchoring junctions in the testis —Sertoli-Sertoli, Sertoli-spermatogonia, Sertoli-spermatocyte and Sertoli-spermatid cell unions [80, 81]— might require complex signaling that helps the progression of cells from base to lumen of the seminiferous tubule during spermatogenesis and subsequent spermiogenesis (Figures 9 and 10). One might think that the long zygotene cilia could be implicated in the signaling pathway between spermatocytes in coordination with Sertoli cells, at least during the first stages of prophase I. Given that ciliopathies present various anomalies in the assembly or regulation of the ciliary and flagellar axoneme [82], more studies will be necessary to analyze the relationship between meiotic cilia and sterility. The present work has described the zygotene cilium in mouse spermatocytes, opening the line of research for future studies to unravel the importance of meiotic ciliogenesis.

## MATERIAL AND METHODS

### Materials

Testes from adult C57BL/6 (wild-type, WT) and genetically modified *Ankrd31*^*–/–*^ [41] male mice were used for this study. All animal procedures were approved by local and regional ethics committees (UAM Ethics Committee for Research and Animal Welfare) and performed according to European Union guidelines.

### Squashing procedure and immunofluorescence microscopy

Seminiferous tubules were fixed and processed following previously described protocols for the squashing technique [83].

Acetylated Tubulin was detected with a mouse monoclonal antibody (Sigma, T7461) at a 1:100 dilution. Tubulin was detected with a mouse monoclonal antibody (Abcam, ab7291) at a 1:100 dilution. Centrin 3 (CETN3) was detected with a mouse monoclonal antibody (Novus, H00001070-M01) at 1:100 dilution. ARL13B detected with a mouse monoclonal antibody (Proteintech, 10002155) at a 1:30 dilution. SYCP3 was detected with either a mouse monoclonal antibody against mouse SYCP3 (Abcam, ab97672) or a rabbit polyclonal antibody recognizing mouse SYCP3 (Abcam, ab15093) both at a 1:50 dilution. SYCP1 was detected with a mouse monoclonal antibody against mouse SYCP1 (Abcam, ab15090) at a 1:50 dilution. TEX14 was detected with a rabbit polyclonal antibody (Proteintech, 18351-1-AP) at a 1:30 dilution. VASA was detected with a rabbit polyclonal against mouse DDX4/MVH (Abcam, ab13840) at a 1:100 dilution. TRF2 was detected with a rabbit polyclonal antibody (Novus, 56321) at a 1:50 dilution. Corresponding secondary antibodies were used against rabbit and mouse IgGs conjugated with either AMCA, Alexa 488, Alexa 555 and Alexa 647 (Molecular Probes), all of them used at a 1:100 dilution.

Immunofluorescence images and stacks were collected on an Olympus BX61 microscope equipped with epifluorescence optics, a motorized Z-drive, and Olympus DP74digital cameras controlled by Cellsens software (Olympus Life Science). Finally, images were processed with ImageJ (National Institute of Health, USA; http://rsb.info.nih.gov/ij) or/and Adobe Photoshop softwares.

### Histology

For histological cryosections, dissected testes were included in OCT (Sakura Finetek Europe) and immediately frozen. The OCT blocks were cut in 10 μm thick sections using a cryostat. The sample slides were allowed to dry at room temperature and hydrated in PBS 3 times followed by fixation with 4% PFA for 10 minutes. Slides were washed in PBS twice and permeabilized in PBS/0.1% Triton X-100 for 20 minutes. Slides were washed in PBS 3 times and blocked with PBS/2% BSA for 30 minutes before being used for immunofluorescence protocol.

### Quantification analysis

The presence of cilia was quantified in 100 zygotene spermatocyte of three biological replicates.

## ACKOWLEDGEMENTS

We express our sincere thanks to Attila Toth for providing the *Ankrd31*^*–/–*^ mice, and to Ignacio Moreno de Alborán and Elisa Carrasco for scientific support. This work was funded by BIOUAM02-2020 to J.P. and R.G.

## AUTHOR CONTRIBUTIONS

RG, FG and PL conceived the study; PL, SPM and IH performed all experiments; PL, SPM, IH, FG, JP and RG analyzed the results; PL, FG and RG wrote the paper with some contributions from the other co-authors.

## CONFLICT OF INTEREST

The authors declare that they have no conflict of interest.

## Supplementary Figures

**Supplementary Figure 1.**
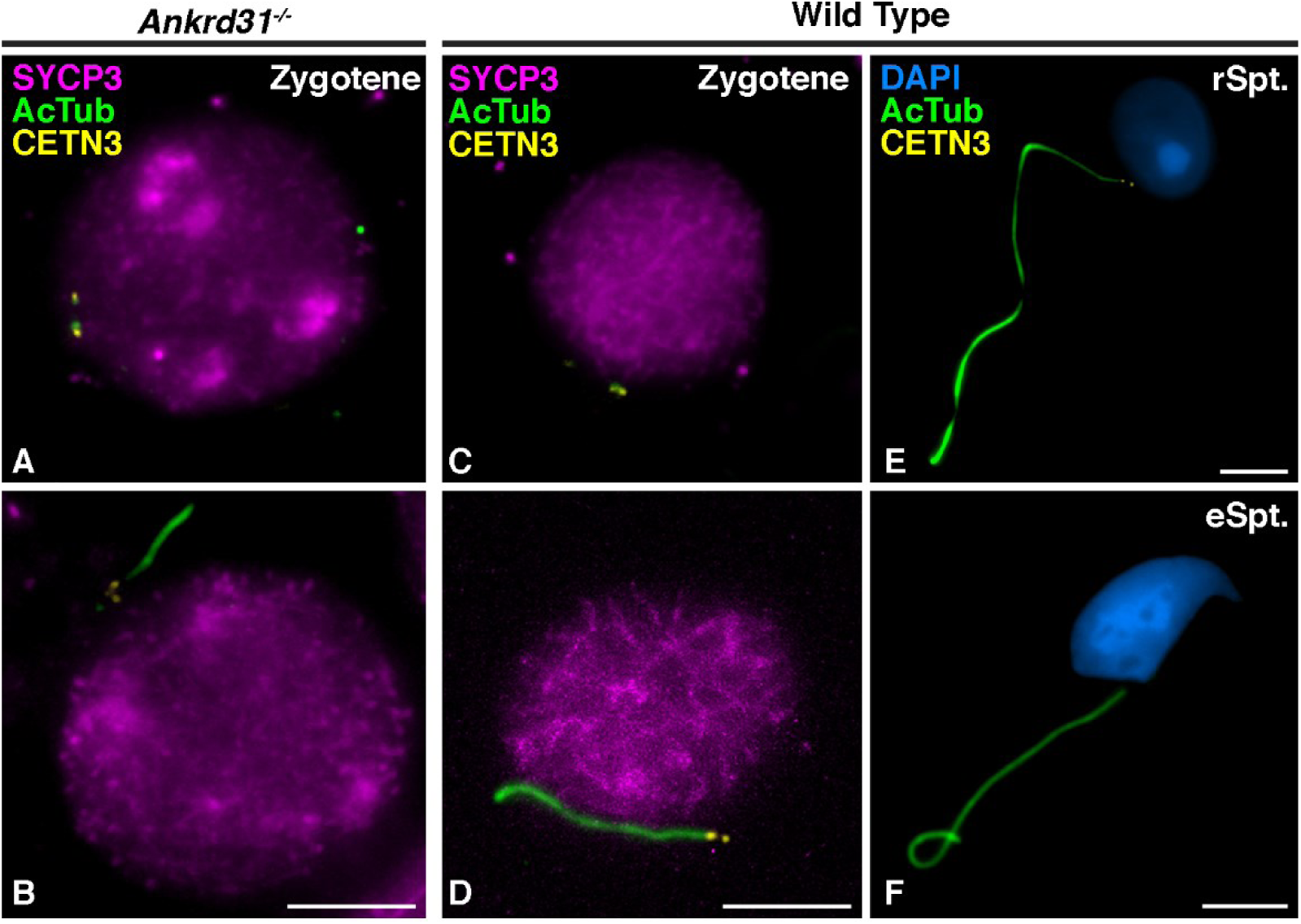
Detection of the cilia in *Ankrd31*^-/-^ sterile mice and WT spermatocytes. Triple immunolabelling of SYCP3 (magenta), acetylated Tubulin (AcTub) (green) and Centrin 3 (CETN3) (yellow) on mouse spermatocytes of *Ankrd3*^*-/-*^ (A,B) at (A) zygotene, and (B) zygotene presenting cilium. Mouse spermatocytes of WT (C,D) at (C) zygotene, and (D) zygotene presenting cilium, € Early round spermatid (rSpt.) and (F) mature elongated spermatid (eSpt.). Scale bar in B, D, E and F represent 5 μm.

**Supplementary Figure 2.**
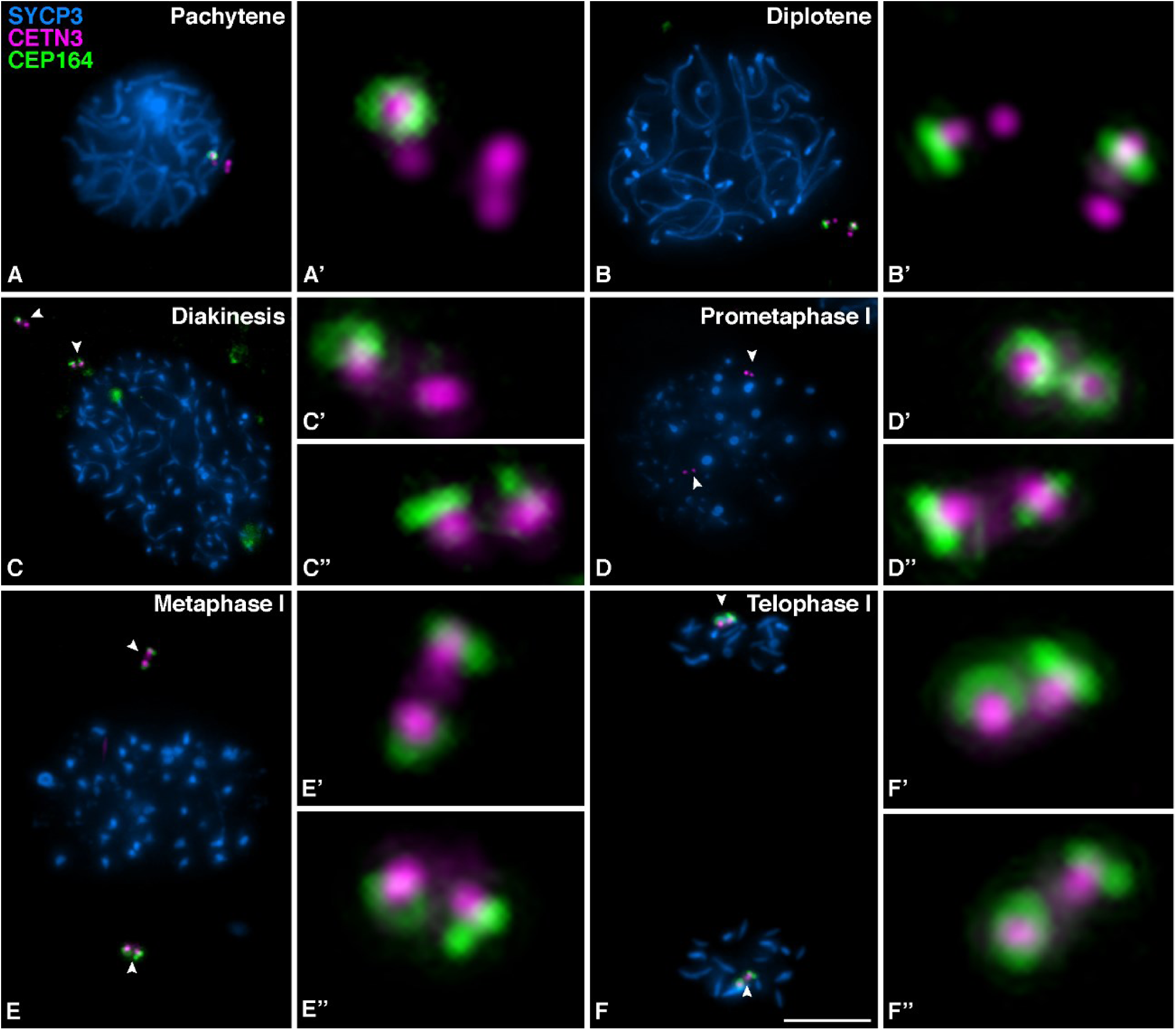
Distribution of CEP164 in mouse spermatocytes during the first meiotic division. Triple immunolabelling of SYCP3 (blue), Centrin 3 (CETN3) (red) and CEP164 (green) in mouse spermatocytes at (A) Pachytene (B) Diplotene, (C) Diakinesis, (D) Prometaphase I, (E) Metaphase I, (F) Telophase I. For images A’-H’ the 300X magnification of the centrosomes is shown (white arrows). Scale bar in H represents 5 μm.

**Supplementary Figure 3.**
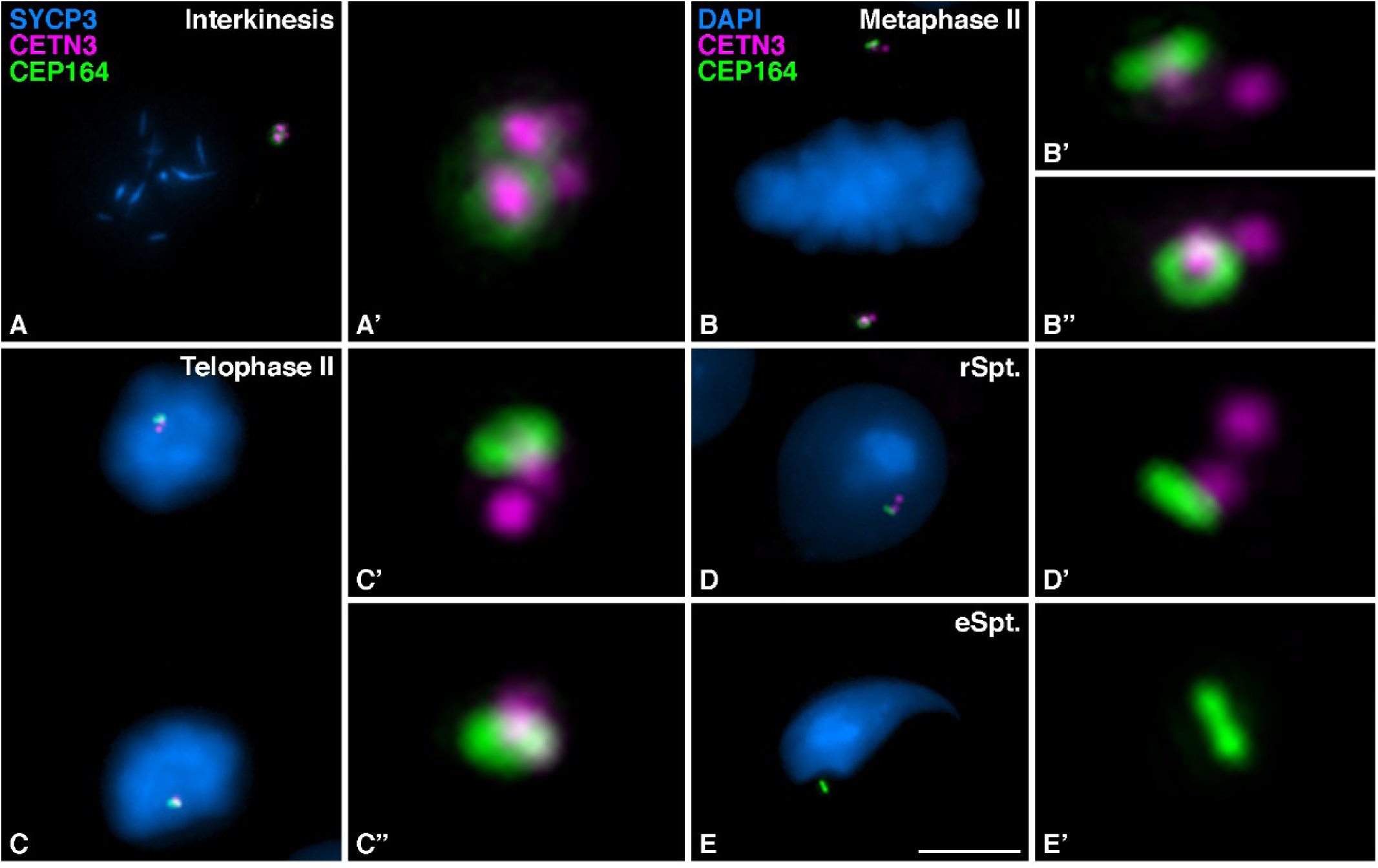
Distribution of CEP164 in mouse spermatocytes during the second meiotic division and spermiogenesis. Triple immunolabelling of SYCP3 (magenta), Centrin 3 (CETN3) (yellow) and CEP164 (green) on squashed WT mouse spermatocytes at (A) Interkinesis. And double immunolabelling of Centrin 3 (CETN3) (yellow) and CEP164 (green), with chromatin stained with DAPI (blue) at (B) Metaphase II, (C) Telophase II (D) Early round spermatid (rSpt.) and mature elongated spermatid (eSpt.). For images A’-E’ the 300X magnification of the centrosomes is shown. Scale bar in E’ represents 5 μm.

## Notes

### Competing Interest Statement

The authors have declared no competing interest.

